# Neutrophil Transcriptomics in SLE: Exploring Intrinsic, Ex Vivo Adaptation, and CAR T-Cell Therapy-Induced Changes

**DOI:** 10.1101/2025.11.03.686294

**Authors:** Ehsan Dehdashtian, Guangnan Hu, Leah Whiteman, Md Tanimul Islam, Stefania Gallucci, Manuel Garber, Dominic Borie, Georg Schett, Roberto Caricchio

## Abstract

**Objectives:** Systemic lupus erythematosus (SLE) is an autoimmune disease characterized by dysregulation of the adaptive and innate immunityThis study aimed to identify transcriptomic differences in neutrophils from SLE patients and healthy individuals, analyze ex vivo adaptation dynamics, and evaluate the impact of chimeric antigen receptor (CAR) T-cell therapy on neutrophil transcriptomic profiles.

**Methods:** Neutrophils were isolated via negative selection from seven SLE patients and three healthy individuals. RNA sequencing was performed to assess transcriptomic differences, ex vivo dynamics over 60 minutes, and responses to lipopolysaccharide (LPS) stimulation. Additionally, longitudinal transcriptomic data from an SLE patient undergoing KYV-101 anti-CD19 CAR T-cell therapy were evaluated.

**Results:** We identified 258 differentially expressed genes (DEGs) consistently distinguishing SLE from healthy neutrophils; they spanned multiple clusters, enriched in interferon-related and DNA damage repair genes (upregulated), and ribosomal protein genes (downregulated). Ex vivo adaptation revealed shared activation pathways, such as NF-κB and apoptosis, in both groups. LPS stimulation highlighted overlapping inflammatory responses, demonstrating retained functional capacities in SLE neutrophils. Following CAR T-cell therapy of an SLE patient,neutrophil transcriptomic profiles realigned with healthy controls by three months post-treatment.

**Conclusions:** Neutrophils in SLE exhibit intrinsic, disease-specific transcriptomic alterations while sharing ex vivo adaptation dynamics with healthy individuals. The disease-specific alterations appear to be modifiable through targeted therapeutic intervention, as anti-CD19 CAR T-cell therapy resets neutrophil gene expression toward healthy patterns despite targeting B cells rather than neutrophils directly. These findings provide insights into SLE pathogenesis and highlight potential therapeutic strategies targeting both adaptive and innate immunity.

## Introduction

Systemic lupus erythematosus (SLE) is a complex autoimmune disorder that disproportionately affects women of reproductive age. It is marked by the production of autoantibodies, immune complex (IC) deposition, and a profound inflammatory response that can affect nearly every organ in the body. (1). Traditionally, SLE pathogenesis is attributed to dysfunctions in the adaptive immune system. However, emerging evidence underscores the role of the innate immune system in both the initiation and progression of the disease (2).

Neutrophils, as key players of the innate immune system, have gained significant attention in recent years for their role in SLE. Neutrophils from SLE patients exhibit dysregulated functions, including enhanced NETosis, accelerated cell death, increased reactive oxygen species (ROS) production, and elevated secretion of pro-inflammatory cytokines such as TNF-α, IFN-α and IFN-γ (2–5). These aberrant neutrophil activities directly contribute to tissue damage, immune complex formation, and sustained inflammatory responses characteristic of SLE pathogenesis (2). Additionally, transcriptomic studies of neutrophils have provided critical insights into SLE pathogenesis, revealing aberrant interferon signatures, dysregulated apoptotic pathways, and enhanced neutrophil activation states that contribute to chronic inflammation and tissue damage (6, 7).

Neutrophils play a critical role in SLE pathogenesis, yet it remains unclear to what extent their dysregulated phenotype is driven by the lupus environment versus intrinsic abnormalities. This study investigated the persistence of disease-specific transcriptional signatures in neutrophils from SLE patients after their removal from the inflammatory milieu. By analyzing ex vivo transcriptional adaptations in neutrophils maintained in stimulant-free conditions, we aim to distinguish intrinsic neutrophil dysfunction from those induced by the SLE inflammatory milieu. Additionally, we examined neutrophil responses to lipopolysaccharide (LPS) stimulation to assess functional differences between SLE and healthy controls.

In recent years, CAR T-cell therapy has demonstrated promising results in patients with SLE who are refractory to conventional treatments (8–10). Since both arms of innate and adaptive immune system are involved in SLE pathogenesis, we aimed to investigate the impact of this therapy on the gene expression profile of neutrophils from an SLE patient undergoing CD19 CAR T-cell therapy.

## Methods

### Patients and Public Involvement

Patients and/or the public were not involved in the design, conduction, reporting, or dissemination of this research.

### Patients and Blood Collection

Seven SLE patients were selected from the UMass Chan Lupus Center (UCLC), a prospective cohort approved by the local IRB. All the patients met at least four criteria of the 2019 EULAR/ACR classification criteria (11). Three healthy, age, sex and race matched controls were also included. Demographic data are shown in Table 1. After informed consent, whole blood was collected in K2 EDTA coated tubes (BD Vacutainer 366643). In addition to the seven patients, a 28-year-old woman with refractory SLE, who underwent lymphodepletion and subsequent CD19 CAR T-cell therapy with KYV-101 (Kyverna Therapeutics, Inc.) via expanded access through a Single-Patient Investigational New Drug application, was also included. Blood samples from this patient were collected at 10 time points: twice before CD19 CAR T-cell infusion and eight times afterwards.

**Table 1.** Demographic and clinical characteristics of study participants. Demographic data, disease characteristics, and medication usage of healthy controls (n=3) and SLE patients (n=7) included in the study. Age, disease duration, and SLEDAI (SLE Disease Activity Index) are presented as median with first (Q1) and third (Q3) quartiles. Medication usage is shown as percentage of SLE patients receiving each treatment. Race/ethnicity is presented as absolute numbers.

|  | Healthy | SLE |
| --- | --- | --- |
| **Age, Median (Q1, Q3) years** | **36 (22, 40)** | **34 (31, 57)** |
| **Sex, % Female** | **100** | **71.4** |
| **Race/ Ethnicity, No.** |  |  |
| **African American** | **1** | **2** |
| **White** | **2** | **3** |
| **Other** | **0** | **2** |
| **Disease Duration, yr. Median (Q1, Q3)** | **-** | **7 (2, 9)** |
| **Medication Usage, %** |  |  |
| **Hydroxychloroquine** | **-** | **86** |
| **Mycophenolate Mofetil** | **-** | **57** |
| **Prednisone** | **-** | **43** |
| **SLEDAI, Median (Q1, Q3)** | **-** | **0 (0, 6)** |

### Neutrophil Purification and Culture

Blood samples were incubated at room temperature for 1 hour and then centrifuged at 1,500 × *g* for 15 minutes. The buffy coat was collected, and neutrophils were isolated by negative selection using the EasySep™ Direct Human Neutrophil Isolation Kit (STEMCELL Technologies, catalog no. 19666), according to the manufacturer’s instructions. For the patient undergoing CAR T-cell therapy, neutrophils were purified directly from whole blood without prior centrifugation or buffy coat separation, using the same kit. This modified approach was implemented due to the limited blood sample volume. Following isolation, purity was confirmed via flow cytometry using CD45, CD16, and CD66b as neutrophil markers (supplementary Fig. 1).

After isolation and cell counting, neutrophils were centrifuged at 400 × *g* for 10 minutes with the acceleration and brake set to 4. The cell pellet was resuspended at 2 × 10^6 cells/mL in RPMI 1640 medium (phenol red–free) supplemented with 1% L-glutamine, then seeded at 1 × 10^6 cells/well in 24-well tissue culture-treated plates (Corning Costar, catalog no. 3526). Neutrophils were either stimulated with LPS (100 ng/mL) or left unstimulated and incubated at 37°C with 5% CO. Unstimulated cells were pelleted at 0, 30, and 60 minutes, whereas LPS-stimulated neutrophils were pelleted at 60 minutes to capture early transcriptional changes while avoiding spontaneous neutrophil activation and NETosis that occurs rapidly in ex vivo conditions (12). For the CAR T-cell therapy patient, neutrophils were pelleted only at time 0, without incubation. All pelleting steps were performed at 400 × *g* for 10 minutes. Pellets were then resuspended in 500 µL Tri-Reagent® (Zymo Research, catalog no. R2050-1-200) and stored at –80°C.

### RNA Extraction and RNA Sequencing Library Preparation

The RNA from neutrophils was extracted using TRI Reagent Solution protocol (ThermoFisher Scientific #AM9738). Purity and integrity were confirmed using the TapeStation system (Agilent Technologies, CA, USA). mRNA was purified using the NEBNext Poly(A) mRNA Magnetic Isolation module (NEB #E7490S/L). After that, libraries were prepared using the xGEN Broad-Range RNA Library Prep Kit (IDT #10009813) and xGen Normalase UDI Primers (IDT # 10009796). The prepared libraries were enzymatically pooled using the xGen Normalase Module (IDT #10009793) and sequenced, generating 150 bp paired-end reads (Azenta Life Sciences, NJ, USA).

### Statistical Analysis

We generated FASTQ files from RNA sequencing and assessed their quality using FastQC. We mapped the reads to the GRCh38 reference human genome using RSEM v1.3.3 to quantify the number of reads per gene. We analyzed the resulting count matrix with the R package DESeq2 (v1.44.0) (13) and identified differentially expressed genes (DEGs) using thresholds of fold change > 1.5 and an adjusted p-value < 0.05. We generated principal component analysis (PCA) plots using the plotPCA function in DESeq2 based on the top 2000 most variable genes, and heatmaps using data normalized with the EdgeR function, log2(CPM + 4). To assess whether disease status (SLE vs. healthy) affected the pattern of time-dependent gene expression changes, we incorporated DESeq2 interaction term analysis. we used the design formula: design = ∼ SLE_Health + Time_point + SLE_Health:Time_point. To assess whether including the interaction term improved the model, we performed a likelihood ratio test (LRT) comparing this full model to a reduced model (design = ∼ SLE_Health + Time_point). Gene expression with an adjusted p-value ≤ 0.05 in this test were considered to be significant based on both time factor and disease status. We identified enriched pathways through hypergeometric testing with the Kyoto Encyclopedia of Genes and Genomes (KEGG) database and defined significantly enriched pathways using a false discovery rate (FDR) cutoff of < 0.05. We conducted data analysis using GraphPad Prism 10 software. We generated heatmaps with the ComplexHeatmap package in R, and created all other plots using either the ggplot2 package in R or GraphPad Prism 10 software.To validate our findings against external datasets, we generated a combined heatmap incorporating data from the GSE139360 dataset by Mistry et al. (6). Gene Set Variation Analysis (GSVA) scores were calculated for each intrinsic DEG cluster to assess functional implications across datasets. Two-way ANOVA tests were performed on the GSVA scores to identify significant variations.

## Results

### Neutrophils from healthy individuals and SLE patients display different intrinsic transcriptomic profiles at baseline which, however, undergo similar changes over time

We conducted a PCA to characterize the overall transcriptomic profiles of neutrophils from healthy and SLE individuals following 0, 30, and 60 minutes of ex vivo culture. The PCA plot shows a distinct separation between healthy and SLE neutrophils primarily along the PC1 axis, highlighting intrinsic differences in their transcriptomic profiles. However, both groups follow similar trajectories over time along the PC2 axis (Fig. 1).

**Fig. 1:**
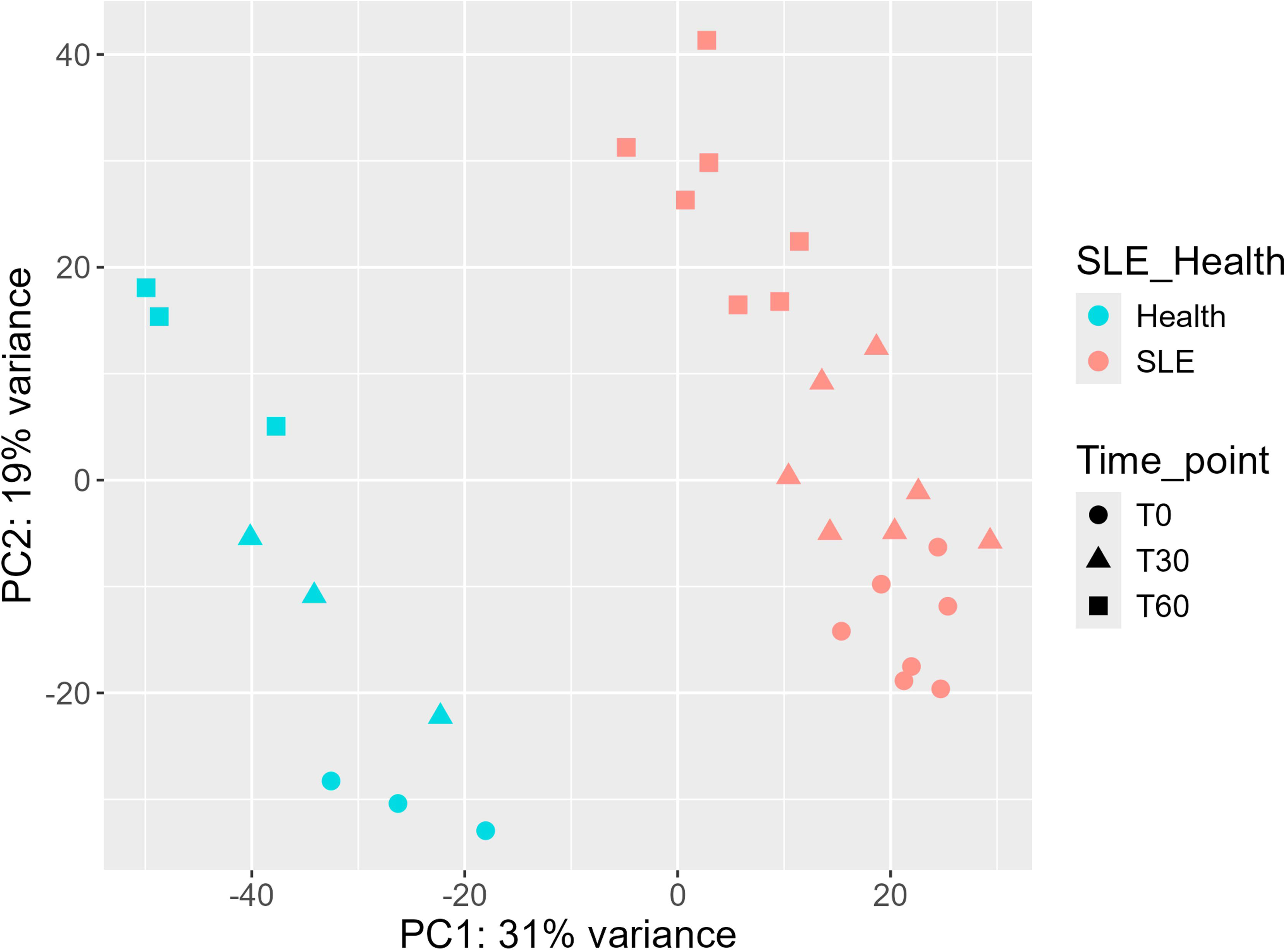
Principal component analysis (PCA). PCA was conducted on neutrophils from healthy and SLE individuals at three time points (0, 30, and 60 minutes). The PCA plot shows a clear separation between healthy and SLE samples along the PC1 axis, indicating intrinsic differences, and a progression along the PC2 axis representing longitudinal transcriptomic shifts.

This parallel progression suggests that, despite inherent differences, neutrophils from healthy and SLE individuals undergo comparable transcriptomic shifts over time. Guided by this observation, we used DESeq2 to identify DEGs, confirming that the time factor had no significant interaction with disease status. These findings indicate that the observed time-dependent changes in gene expression are independent of SLE, with both groups displaying similar patterns of transcriptional adaptation during ex vivo culture.

### Impact of Time Progression on Transcriptomic Profiles of Neutrophils

We performed differential gene expression analyses on neutrophils from both healthy individuals and those with SLE at three time points (0, 30, and 60 minutes) following ex vivo culture. Volcano plots (Supplementary Fig. 2A) illustrate the DEGs for each pairwise comparison in both groups. These plots highlight genes that were significantly upregulated or downregulated over time in neutrophils from both healthy and SLE donors.

Venn diagrams further clarify the relationships among DEGs across the three time-point comparisons (Supplementary Fig. 2B). In the first Venn diagram, each set (circle) represents the following comparisons of DEGs combining both healthy and SLE samples: (1) 0 vs. 30, (2) 0 vs. 60 minutes, and (3) 30 vs. 60 minutes. The second and third Venn diagrams show these ex vivo time-dependent DEG patterns separately for healthy controls and SLE patients. While these latter diagrams appear to show different numbers of DEGs between healthy and SLE neutrophils over time, our statistical analysis revealed that, despite having different baseline transcriptional profiles, neutrophils from both groups undergo similar ex vivo adaptations in gene expression over time (except for the three genes of BTG2, SEPTIN4, and GSTO2).

We used heatmaps to visualize the DEGs that are common across groups (Fig. 2A). Based on these heatmaps and Venn diagrams, we observed three main groups of gene expression changes: (1) genes that downregualate significantly from 0 to 30 minutes and continue in the same direction through 60 minutes, (2) genes that upregulate significantly from 0 to 30 minutes and continue in the same direction through 60 minutes, and (3) genes that mainly show significant changes in expression only between 30 and 60 minutes (Fig. 2B).

**Fig. 2:**
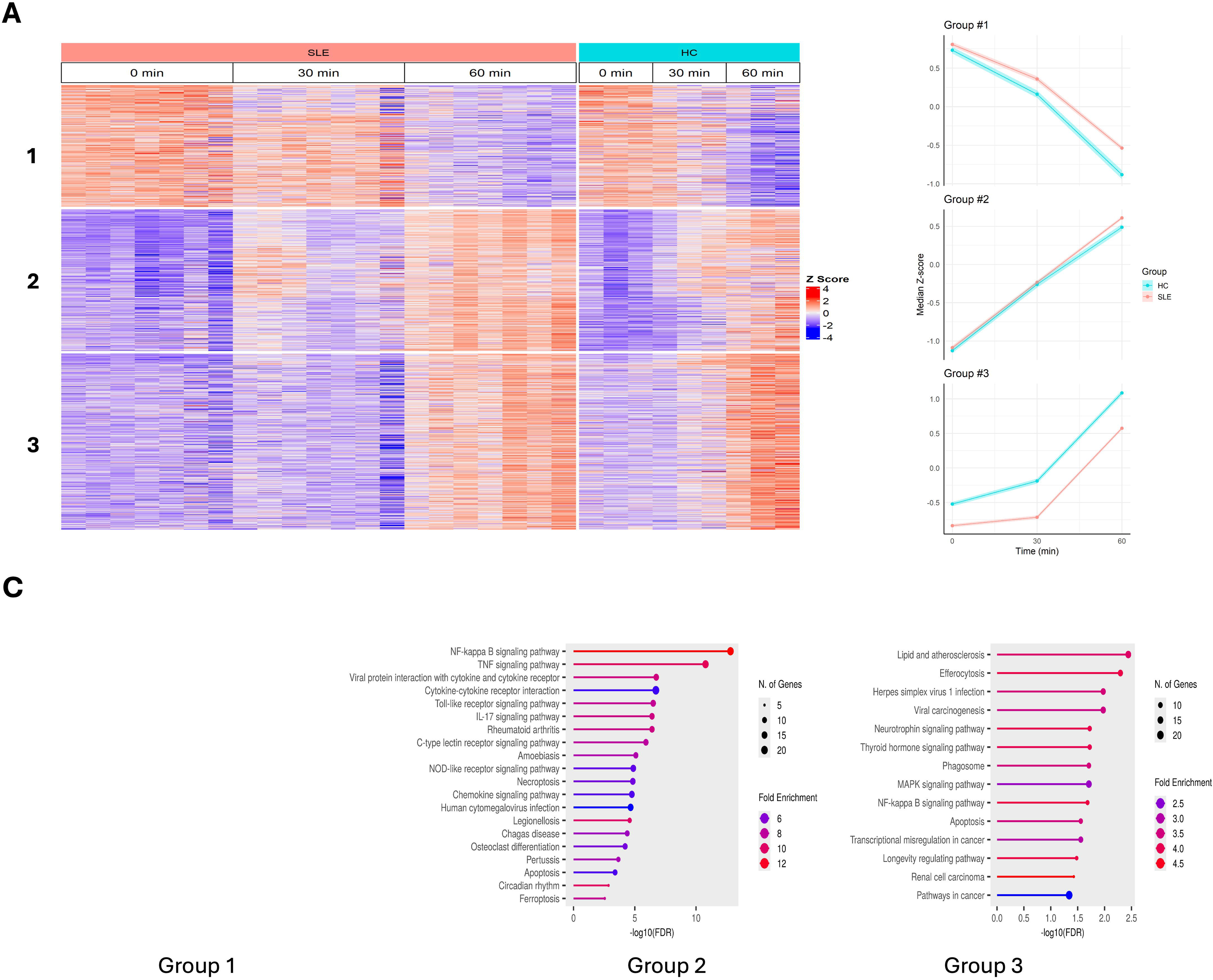
Heatmap of temporal transcriptomics in neutrophils and pathway enrichment analysis. (**A**) Heatmap displaying temporal transcriptomics of neutrophils categorized into three groups:(1) sustained downregulation from time 0 to 60 minutes, (2) sustained upregulation from time 0 to 60 minutes, and (3) changes only between 30 and 60 minutes. **(B)** Graphic Conceptualization of the 3 groups. **(C)** KEGG pathway enrichment analysis identifies key pathways impacted by time progression, including NF-κB activation, TNF signaling, and apoptosis, in neutrophils from both groups.

To identify the biological processes affected by these changes, we performed KEGG pathway enrichment analysis on DEGs from each comparison group (Fig. 2C). The first group showed no enrichment. The second group showed enrichment in multiple inflammatory pathways including NF-κB, TNF, IL-17 signaling and apoptosis. In the third group, while NF-κB signaling persisted, we observed a shift toward cell death-related processes, with significant enrichment in apoptosis, MAPK signaling and longevity regulating pathways. A common feature is that neutrophils undergo activation and cell death over time following isolation as indicated by the consistent activation of the NF-κB and apoptosis pathways (14). This time-dependent progression from broad inflammatory response to focused activation and ultimately cell death pathways provides insights into neutrophil behavior under ex vivo conditions, though we acknowledge these changes may partly reflect artificial activation due to isolation procedures and culture conditions.

### Intrinsic Differences in Neutrophil Transcriptomics of Healthy and SLE Individuals

A Venn diagram was generated to illustrate the DEG patterns between neutrophils from healthy and SLE individuals across three time points: 0, 30, and 60 minutes (Supplementary Fig. 3). The DEGs were identified using the thresholds of fold change > 2 and an adjusted p-value < 0.01. This diagram reveals the overlapping and unique gene sets for each comparison. Notably, 262 DEGs were identified as common across all three time points, suggesting stable intrinsic differences in transcriptomic profiles between neutrophils from healthy and SLE individuals, regardless of ex vivo culture duration. Out of these 262 DEGs, four genes were related to the Y chromosome and were excluded from future analyses, and the rest are referred to as intrinsic DEGs.

To assess whether these intrinsic differences vary with disease activity, we examined their relationship with clinical parameters. Disease activity was measured using SLEDAI score for all patients. When patients were stratified into groups based on their SLEDAI scores, comparative analysis revealed no significant differences in neutrophil gene expression patterns between these groups (not shown). Similarly, when patients were grouped according to their medication regimens, no significant differences in transcriptomic profiles were observed between treatment groups (not shown). These findings further support that the observed transcriptomic differences reflect intrinsic disease-specific characteristics that remain stable regardless of clinical disease activity or therapeutic intervention.

We categorized the intrinsic DEGs into several clusters based on information from KEGG, Reactome, WikiPathways, Hallmark, and GO databases. Our first five clusters include genes related to interferon (IFN) signaling, Rho GTPase activity, cytoskeleton organization, DNA damage repair, and ribosome function. We formed a sixth cluster from long non-coding RNAs (lncRNAs) and grouped the less well-characterized DEGs into a final cluster labeled “other.” We generated heatmaps to display the expression patterns of these gene clusters across the three time points (Fig. 3).

**Fig. 3:**
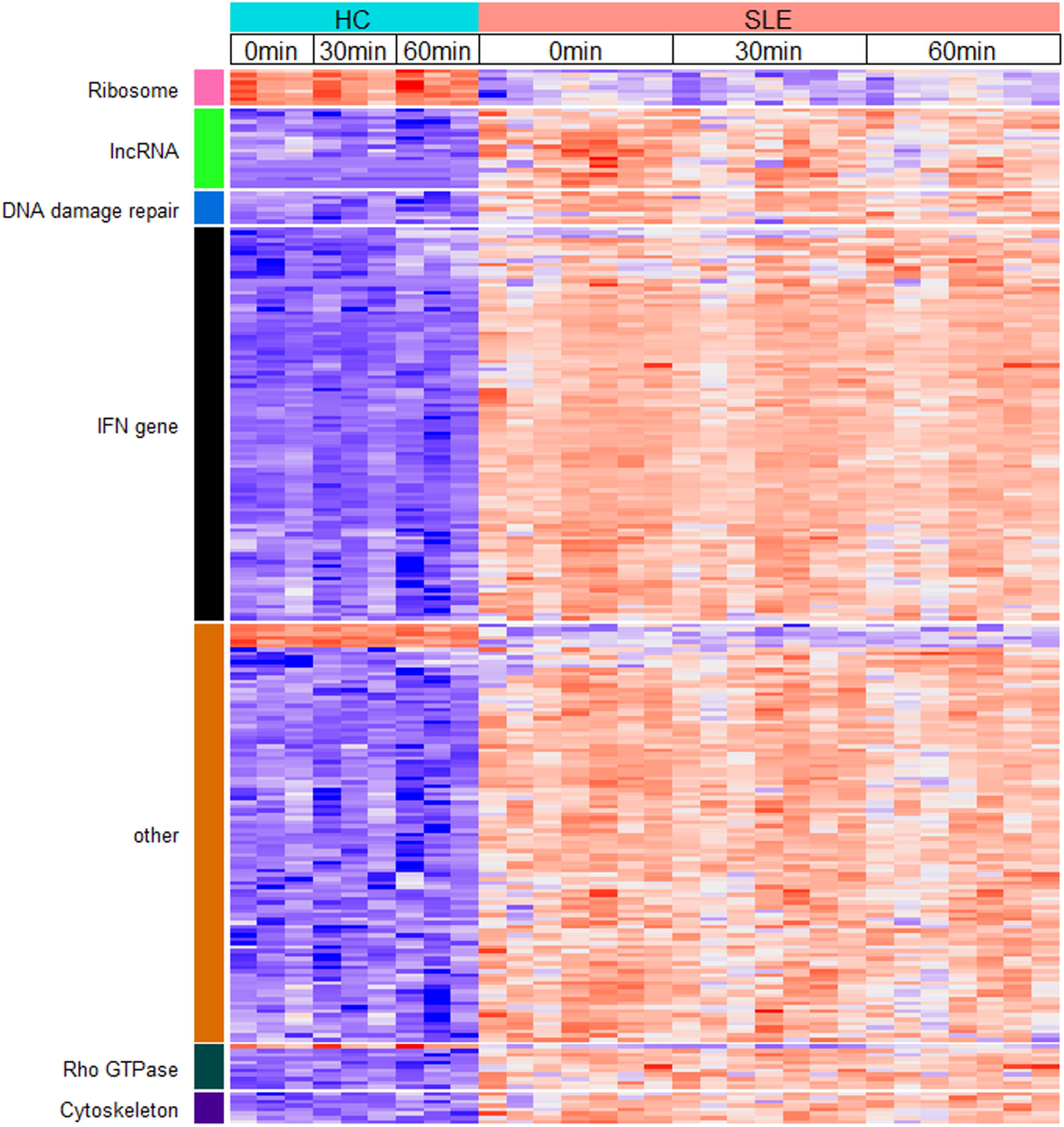
Clustered heatmap of intrinsic DEGs. Heatmap showing expression patterns of intrinsic DEGs categorized into seven clusters including interferon-related, Rho GTPase, cytoskeleton, DNA damage repair, ribosome, and long non-coding RNA (lncRNA). The “other” cluster includes genes with no clear pathway enrichment.

The first cluster (IFN-related genes) shows significant upregulation in SLE patients compared to healthy individuals. The second and third clusters contain genes from the Rho GTPase and cytoskeleton gene sets, which regulate intracellular trafficking, cell migration, cell polarity, gene expression, and cell proliferation. These biological processes underpin essential immune functions such as cellular trafficking, phagocytosis, and antigen recognition (15, 16). The DNA damage repair gene set, which forms the next cluster, also exhibits upregulation in SLE samples, while the ribosome gene cluster shows downregulation in SLE neutrophils relative to healthy controls. Additionally, our lncRNA cluster—which includes, among others, JPX, FTX, NRIR, LINC02574, and IRF1-AS1—displays higher expression in SLE samples. The “other” cluster contains DEGs upregulated in SLE neutrophils that lack significant enrichment information in the databases we used. Based on literature review, we further categorized several well-characterized genes from the “other” cluster into six groups (Supplementary Fig. 4 and Supplementary Table 1).

Comparison with an independent dataset (Mistry et al.) (6) confirmed that the intrinsic transcriptomic differences we identified between healthy and SLE neutrophils are consistent with previous findings (Supplementary Fig. 5). Additionally, analysis of pathway activity through GSVA scores (Fig. 4) revealed significant variations in functional pathways between healthy and SLE neutrophils. Notably, the IFN-related genes, along with other intrinsic gene clusters, maintained their differential expression patterns in SLE neutrophils throughout the culture adaptation period, suggesting these transcriptomic alterations are long-lived and intrinsic to SLE neutrophils rather than simply reflecting their immediate inflammatory milieu.

**Fig. 4.**
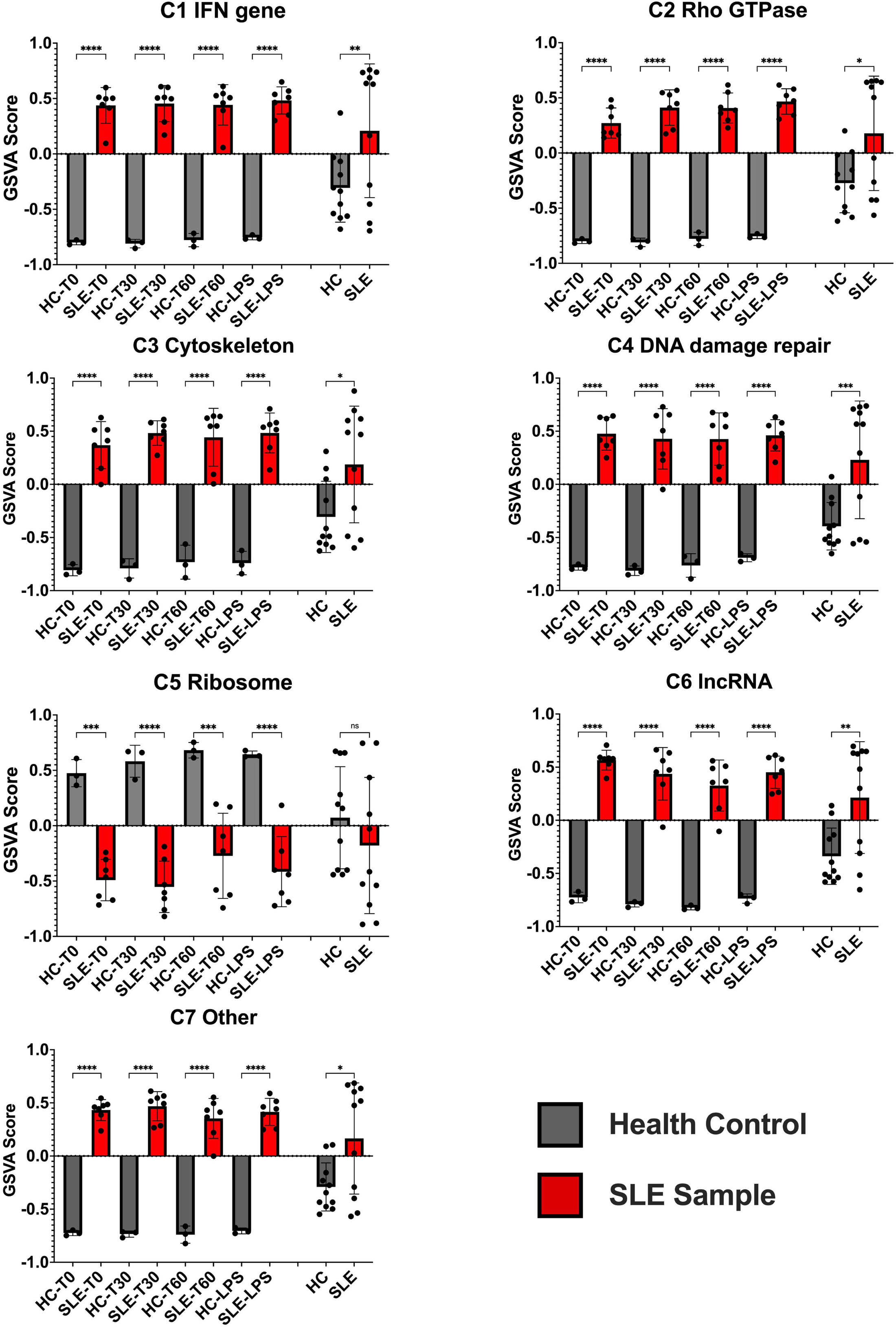
GSVA scores of intrinsic DEG clusters across datasets. GSVA scores calculated for each intrinsic DEG cluster, comparing pathway activities in neutrophils from healthy individuals and SLE patients. Data from this study are displayed alongside corresponding GSVA scores from the GSE139360 dataset. Two-way ANOVA tests were used to compare the scores, revealing significant differences in clusters associated with intrinsic transcriptomic alterations in SLE neutrophils.

### Distinct DEGs in SLE Neutrophils: Cholesterol Handling and Immune Function

Apart from the enrichment tests, some of the differences in intrinsic DEGs between healthy and SLE neutrophils are of particular interest. Among those ABCA1, MDK and GRAMD1B which are involved in cholesterol metabolism, are worth mentioning. GRAMD1B is an IFN-I inducible gene that codes for a protein involved in the transport of HDL and LDL-derived cholesterol from the plasma membrane to the endoplasmic reticulum (17, 18). Conversely, ABCA1 is an integral plasma membrane protein that plays a major role in exporting excess cholesterol out of the cell (19). MDK is involved in decreasing ABCA1 expression, thus inhibiting the cholesterol efflux out of cell (20). While all these genes are upregulated in neutrophils from SLE patients compared to healthy individuals, ABCA1 shows a markedly lower fold change than the other two genes, suggesting potential abnormalities in cholesterol metabolism within SLE patients’ neutrophils (Supplementary Fig. 6). Further evidence suggesting altered cholesterol metabolism in SLE patient neutrophils is the upregulation of C7orf50, which encodes the hormone Cholesin that functions to inhibit cholesterol synthesis in the liver (21).

Another gene of interest from the “other” cluster is TNFSF13B, which encodes B cell activating factor (BAFF), a key player in SLE pathogenesis (22). Quantitative analysis revealed significant upregulation of TNFSF13B (more than 2-fold) in neutrophils from SLE patients compared to healthy individuals. Notably, this upregulation was consistently maintained across all three time points examined, suggesting it represents a stable intrinsic feature of SLE neutrophils. The upregulation of BAFF in neutrophils may represent a critical mechanism by which these cells contribute to B cell dysregulation and autoantibody production in SLE pathogenesis (4, 23).

MICB is another upregulated gene in neutrophils of SLE patients. It is upregulated under cellular stress and DNA damage and can tag cells for elimination by natural killer (NK) cells (24). This suggests that neutrophil elimination and the release of their cellular contents may continuously occur in SLE. Among the IFN-related genes cluster, MOV10, which can act as an autoantigen in SLE (25), is upregulated in neutrophils of SLE patients compared to healthy individuals.

### LPS Induces a Common Inflammatory Response in Neutrophils While Preserving SLE-Specific Transcriptional Differences

Given the elevated interferon signature in SLE neutrophils, we hypothesized that these cells might exhibit an altered response to pathogen-associated molecular patterns compared to healthy controls. To test this hypothesis, neutrophils from both healthy donors and SLE patients were stimulated with LPS, followed by transcriptomic analysis to detect LPS-induced gene expression changes. Surprisingly, the analysis revealed that both groups shared a common set of DEGs in response to LPS, including classic inflammatory genes such as *IL1B, CXCL2, TNF,* and *CCL20*.(26–29). To visualize these transcriptional changes, we generated a heatmap incorporating both the LPS-induced DEGs and the SLE-specific DEGs, illustrating the common inflammatory response as well as the persistence of disease-specific transcriptional signatures (Fig. 5A).

**Fig. 5:**
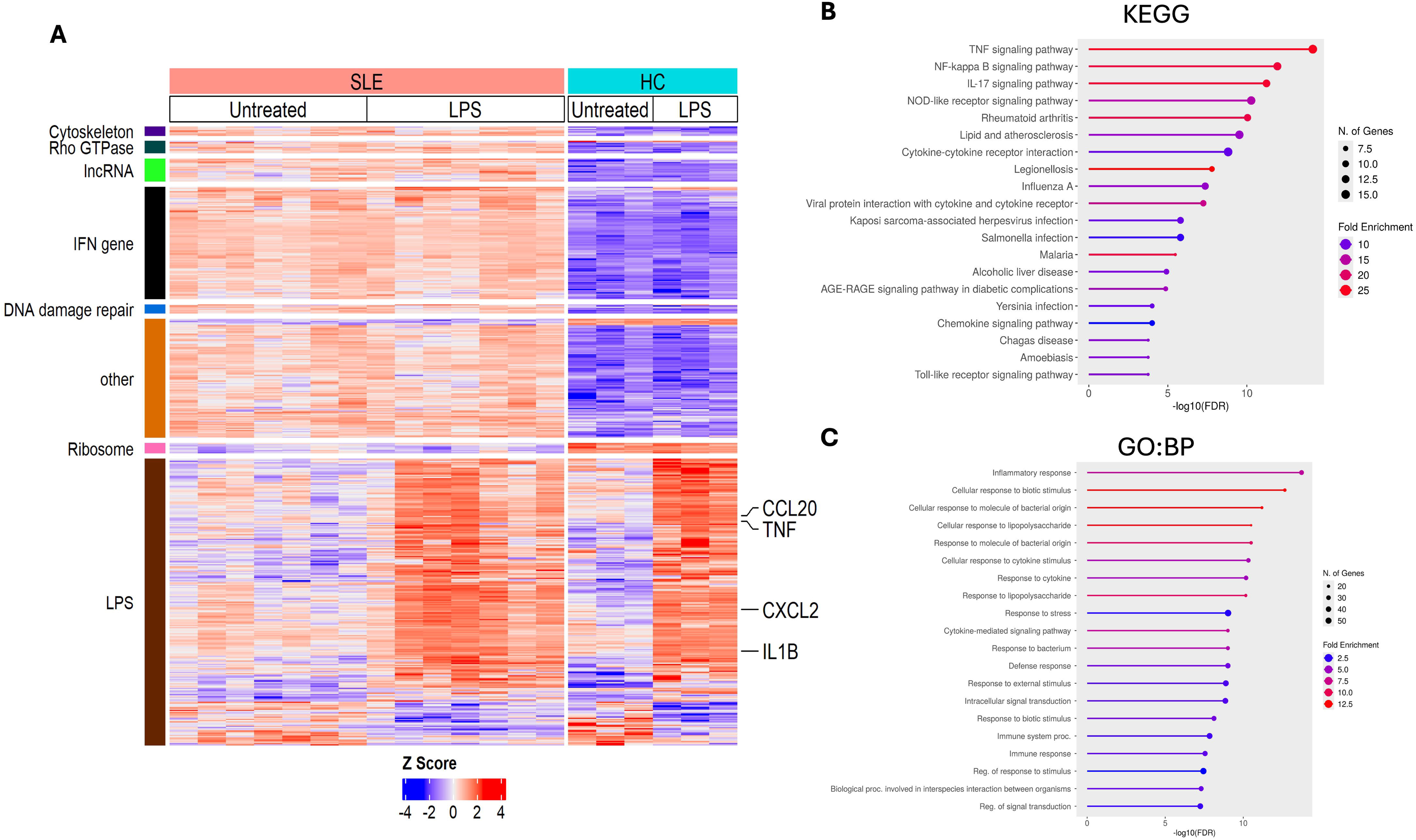
Impact of LPS stimulation on neutrophil transcriptomics. (A) Heatmap displaying shared inflammatory response genes induced by LPS stimulation in neutrophils from both healthy and SLE individuals (labeled as “LPS”), along with the persistence of intrinsic transcriptional differences between neutrophils from both groups (organized in clusters: cytoskeleton, Rho GTPase, lncRNA, IFN genes, DNA damage repair, other, and ribosome) despite LPS treatment. (B) KEGG pathway analysis of LPS-induced DEGs, showing enrichment of key inflammatory pathways such as TNF and NF-_κ_B signaling. (C) GO biological process (GO:BP) analysis revealing significant enrichment of inflammatory and immune response pathways following LPS stimulation.

To further investigate the pathways activated by LPS, we performed KEGG and Gene Ontology (GO) enrichment analyses on the DEGs identified in both healthy and SLE neutrophils in response to LPS (Fig. 5B and 5C). KEGG pathway analysis revealed significant enrichment of inflammatory and immune-related pathways, including TNF signaling, NF-κB activation, and IL-17 signaling, consistent with a robust inflammatory response to LPS. Similarly, GO biological process (GO:BP) analysis highlighted inflammatory response, cellular response to biotic stimulus and LPS as key processes induced by LPS in both groups. These findings indicate that LPS elicits a shared pro-inflammatory transcriptional program in neutrophils, regardless of disease status.

Despite this common response to LPS, we observed that the intrinsic transcriptional differences that distinguish SLE from healthy neutrophils remained intact with minimal response to LPS, further supporting the idea that lupus-associated transcriptional alterations persist even in the presence of external inflammatory stimuli.

### Longitudinal Impact of CAR T-Cell Therapy on Intrinsic Transcriptomic Profiles in SLE Patient Neutrophils

To investigate the longitudinal impact of CD19 CAR T-cell therapy on intrinsic DEGs in neutrophils, we analyzed data from an SLE patient undergoing this treatment. The patient, who had a SLEDAI score of 17 before treatment, received CD19 CAR T-cell therapy in 2024 (day 0). Notably, by 4 months post-treatment, the patient’s SLEDAI score had decreased to 4, indicating significant clinical improvement. Moreover, by month 5 post-infusion the patient was off any immune suppressive therapy.

A heatmap was generated to visualize the expression patterns of the previously identified intrinsic DEGs in the patient’s neutrophils at baseline (pre-treatment) and multiple time points post-treatment: day 3, day 7, day 14, day 21, day 28, day 56, month 3, and month 5. For reference, neutrophil samples from three healthy individuals were included, allowing for a direct comparison between healthy controls and the patient at various stages post-treatment (Fig. 6A).

**Fig. 6:**
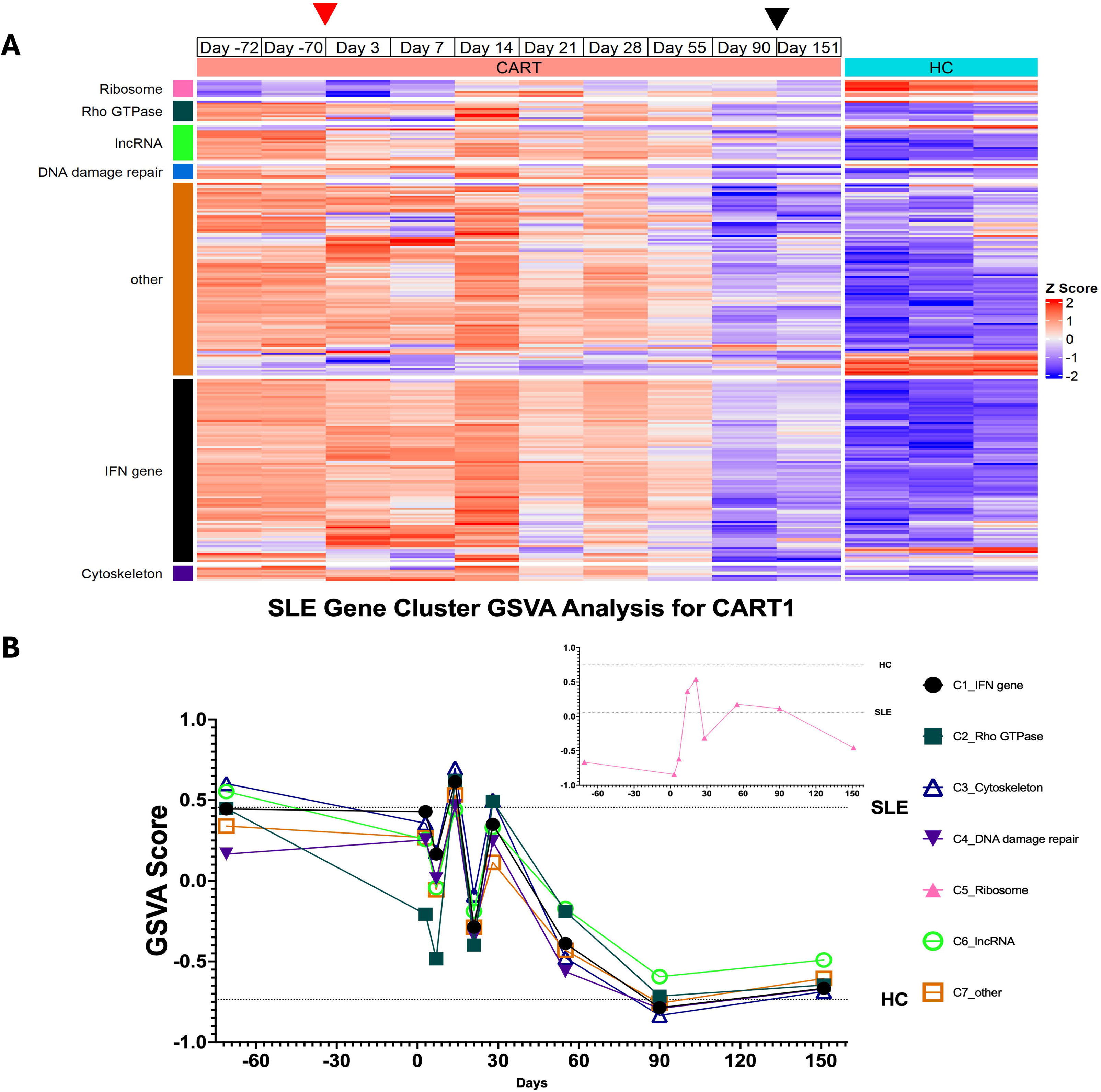
Longitudinal transcriptomic and GSVA score analysis post-CAR-T cell therapy. (A) Heatmap visualizing intrinsic DEG expression in an SLE patient undergoing CAR T-cell therapy across baseline (two pre-treatment time points) and eight post-treatment time points. Samples from three healthy individuals serve as controls. Red arrowhead indicates day 0 (CAR T-cell infusion) and black arrowhead marks when the patient discontinued all immunosuppressive medications. (B) GSVA plots showing pathway activity for intrinsic DEG clusters across time points in the CAR T-cell therapy patient. The analysis reveals dynamic transcriptomic reprogramming, with most clusters approaching healthy transcriptomic profiles by three months post-treatment.

The heatmap clusters genes according to the same pathway-based enrichment categories identified earlier in our analysis, providing insight into how those intrinsic DEGs evolve over time in response to CD19 CAR T-cell therapy. At baseline, the patient’s neutrophil transcriptome differed markedly from healthy controls, with upregulation of interferon, DNA damage repair, Rho GTPase, cytoskeleton, lncRNA, and other gene clusters, while ribosomal structural protein genes were downregulated. Following therapy, we observed modest changes in gene expression during the first two months, but a substantial transition began around 3 months, with the neutrophil expression profile at month 3 and month 5 closely resembling that of healthy controls across most gene clusters.

To provide a more quantitative assessment of these longitudinal changes, GSVA scores were calculated for each time point in the CD19 CAR T-cell therapy patient, based on the same gene clusters used in previous analyses (Fig. 6B). The GSVA plot provides a quantitative view of pathway activity changes over time. As shown in the figure, following initial CAR T-cell therapy, we observed some fluctuation in pathway scores during the first month, followed by a consistent trend toward healthy control values across most gene clusters by month 3. This gradual normalization timeline is consistent with a progressive immune reset process. Except for the ribosome cluster, the gene expression levels in all other clusters resembled those of healthy controls by 3 months post-therapy.

## Discussion

This study provides a comprehensive investigation into the transcriptomic landscape of neutrophils in SLE, offering several novel insights that extend beyond previous work in this field.

By employing a refined neutrophil isolation technique and examining ex vivo transcriptomic adaptations over time, we have introduced an approach that, to the best of our knowledge, has not been previously utilized in SLE neutrophil research. While previous studies have examined neutrophil transcriptomics in SLE (6, 7), our work presents three key innovations: (1) the assessment of transcriptomic stability during ex vivo adaptation, allowing discrimination between intrinsic versus micro-environment induced differences; (2) the evaluation of functional responses to LPS stimulation alongside disease-specific signatures; and (3) the first longitudinal analysis of neutrophil transcriptomic changes following CD19 CAR T-cell therapy in an SLE patient.

Our findings revealed that neutrophils from both healthy and SLE groups exhibited remarkably similar and rapid transcriptional adaptation patterns during culture. This parallel behavior in response to ex vivo conditions suggests that despite disease status and ongoing therapy, SLE neutrophils retain many core functional capabilities similar to healthy controls. Both populations showed coordinated activation of key pathways including NF-κB signaling, inflammatory responses, and regulated progression toward apoptosis. While these shared responses demonstrate the preservation of essential neutrophil functions in SLE despite disease-specific differences, they also highlight the challenges of maintaining neutrophil stability ex vivo and underscore the importance of rapid processing in transcriptomic studies. (30).

Our findings also revealed that while healthy and SLE neutrophils can exhibit similar transcriptomic changes in response to time, there are intrinsic DEGs between the two groups that remain consistently different regardless of the in vitro culture duration. These persistent DEGs likely reflect fundamental differences in the baseline biology of neutrophils from healthy and SLE individuals and possibly also reflect the disease vs health status irrespective of the disease activity. These genes were categorized into several clusters, each providing unique insights into SLE pathogenesis. The upregulation of interferon-related genes in SLE neutrophils aligns with the established role of interferons in SLE pathogenesis (31). This finding reinforces the critical importance of interferon signaling in the molecular mechanism of the disease.

Apart from interferon-related genes, differences in other functional gene clusters were observed between neutrophils from SLE patients and controls. The upregulation of most of Rho GTPase and cytoskeleton gene sets in SLE neutrophils may be due to the aberrant neutrophil activation in SLE patients (32). Increased expression of DNA damage repair genes in neutrophils from SLE patients may reflect efforts by the cells to repair DNA damage caused by various factors, such as elevated levels of reactive oxygen species (ROS) (32–34). The downregulation of ribosomal structural protein gene expression may result from the metabolic reprogramming of cells under immunological stress, which redirects their transcriptomic machinery toward the production of inflammatory genes rather than housekeeping ones (35–37).

JPX and FTX are two of the lncRNAs involved in X chromosome inactivation by upregulating the expression of Xist (38). Despite the increased expression of JPX and FTX in our dataset, the level of Xist does not show a significant difference between neutrophils from healthy individuals and those with SLE. However, Tasl (categorized in “other” cluster), an X-linked member of the TLR7-9 pathway, is consistently upregulated in SLE neutrophils compared to healthy individuals suggesting abnormal X chromosome inactivation in neutrophils of SLE patients similar to their T cells and B cells, as shown by previous studies (39–41). NRIR, LINC02574 and IRF1-AS1 are also among the lncRNA cluster and are considered interferon stimulated genes which are expected to be upregulated in SLE (31, 42–44). Interestingly, IRF1-AS1 depletion in HeLa cells resulted in increased expression of Xist (44). Thus, the overexpression of this gene in neutrophils from SLE patients might contribute to altered X chromosome inactivation and may explain why Xist is not upregulated despite the increased expression of JPX and FTX lncRNAs.

Comparing our intrinsic DEGs with the previous study by Mistry et al. (6) (Fig. 4 & supplementary Fig. 5) confirms the presence of most of the intrinsic DEGs between healthy and patient neutrophils in their dataset as well. However, the GSVA score difference between healthy and SLE individuals regarding intrinsic DEGs clusters is less in their dataset, which may be attributed to the neutrophil purification method using density gradient medium, which inadvertently results in more neutrophil activation compared to using negative selection by magnetic beads (30). The GSVA scores of Ribosome cluster in Mistry et al. datasets didn’t show significant difference between healthy and SLE individuals, possibly due to patients heterogenicity.

Correlation analysis between GSVA scores for our intrinsic DEG clusters and previously identified ISG (45) GSVA scores in SLE samples revealed a fundamental distinction between neutrophils and other immune cells (Supplementary Fig. 7). While the IFN-related cluster (C1) showed significant ISG correlation in neutrophils, clusters C2-C6-representing Rho GTPase, ribosome, cytoskeleton, DNA damage repair, and lncRNA-showed no significant ISG correlation in our neutrophil dataset. In contrast, these same clusters exhibited highly significant ISG correlations across PBMCs (GSE50772) (46), whole blood (SRP136102) (47), and B cells (SRP156583) (48). This IFN-independent pattern of dysregulation in multiple functional pathways distinguishes neutrophil contributions to SLE pathogenesis from the predominantly IFN-driven dysfunction observed in other immune cells, suggesting unique neutrophil-specific mechanisms.

Our LPS stimulation experiment revealed that both healthy and SLE neutrophils demonstrated overlapping inflammatory responses, as evidenced by the upregulation of common inflammatory genes. Importantly, while LPS induced substantial transcriptional changes in both groups, the intrinsic disease-specific signatures that distinguish SLE neutrophils remained intact. This indicates that the mechanisms controlling the SLE-specific gene expression operate independently from pathways responding to infectious stimuli. The coexistence of these parallel inflammatory mechanisms could potentially lead to enhanced tissue damage during infectious challenges, which may help explain the increased risk of complications that SLE patients face during infections (49).

The longitudinal analysis of an SLE patient undergoing CD19 CAR T-cell therapy provided unique insights into neutrophil transcriptomic recovery. The gene expression patterns did not immediately normalize after treatment, indicating that neutrophil transcriptomes may require some time to realign with healthy profiles, highlighting the importance of lupus epigenetics (6, 50). By three months post-therapy, most gene clusters approached healthy neutrophil characteristics, suggesting gradual immunological reset. Notably, this therapeutic approach highlights the interconnectedness of adaptive and innate immune dysfunction in SLE, as evidenced by changes in neutrophil gene expression following CD19-targeted intervention. This study, while only in a single patient, also reveals how anti-CD19 CAR T-cell therapy can drive durable remission through deep B-cell depletion to broadly reset the immune system to patterns seen in healthy individuals.

Our study introduces several methodological advances. By utilizing negative selection for neutrophil isolation, we minimized potential activation artifacts common in traditional isolation techniques. This approach also enabled the uniform isolation of all neutrophils, regardless of their density. Furthermore, the examination of transcriptomic patterns during ex vivo adaptation, complemented by LPS stimulation experiments, offers a more comprehensive and dynamic understanding of neutrophil behavior in SLE.

While our study offers valuable insights, several limitations should be acknowledged. The relatively small sample size and focus on a single CD19 CAR T-cell therapy patient suggest the need for larger and more comprehensive studies. Future research could explore the functional implications of the identified transcriptomic differences and investigate potential therapeutic interventions targeting these molecular pathways.

## Conclusion

This comprehensive transcriptomic analysis of neutrophils in SLE unveils complex molecular alterations that contribute to our understanding of the disease’s pathogenesis. By highlighting intrinsic differences, time-dependent shifts, and inflammatory responses, our study provides a foundation for a deeper understanding of neutrophil involvement in SLE pathogenesis and highlights potential avenues for therapeutic intervention. Finally, our longitudinal analysis of neutrophil transcriptomes following anti-CD19 CAR T-cell therapy, though limited to a single patient, provides compelling evidence of immune reset beyond the targeted B-cell compartment, with neutrophil gene expression profiles normalizing toward healthy patterns by three months post-treatment. This finding suggests that targeted immunotherapy can induce systemic immune reporogramming and restore normal gene expression in innate immune cells, offering a potential explanation for the durable remissions observed with CAR T-cell therapy in SLE.

## Competing Interest

All authors declare they have no competing interests.

## Acknowledgements

We thank all of the families, patients, parents, and carers who contributed to the study and allowed us to use samples and data for this work.

## Contributors

ED contributed to methodology, investigation, analysis, data curation, and manuscript drafting. GH and MTI contributed to methodology, investigation, and data curation. LW contributed to data curation and manuscript editing. SG, DB, MG, and GS reviewed and edited the manuscript. SG and RC supervised the study. RC conceptualized the project, secured funding, and oversaw the project. All authors reviewed and approved the final manuscript for submission.

## Funding

None

## Patient Consent for Publication

Not applicaple.

## Ethics Approval

Written informed consent was obtained from all participants with approval from research ethics committee of institutional review board at UMass Chan Medical School.

## Declaration of generative AI and AI-assisted technologies in the writing process

During the preparation of this work, the authors used Claude ai in order to improve the text and the readability of the text. After using this tool/service, the authors reviewed and edited the content as needed and take(s) full responsibility for the content of the published article.

## Notes

### Competing Interest Statement

The authors have declared no competing interest.

