## Supplementary figures and images for "Neutrophil Transcriptomics in SLE: Exploring Intrinsic, Ex Vivo Adaptation, and CAR T-Cell Therapy-Induced Changes"

### supplementary Fig. 1

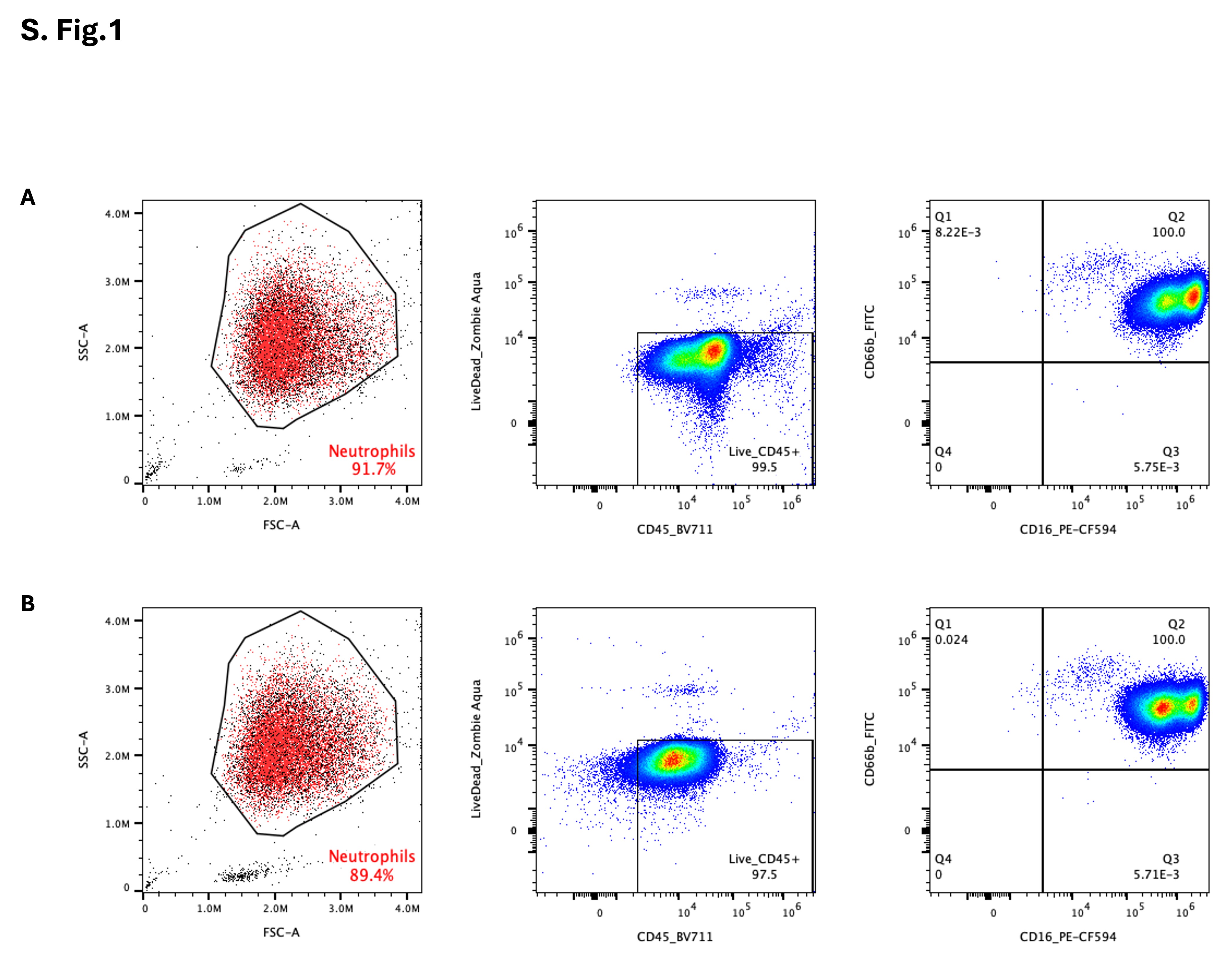

### supplementary Fig. 2

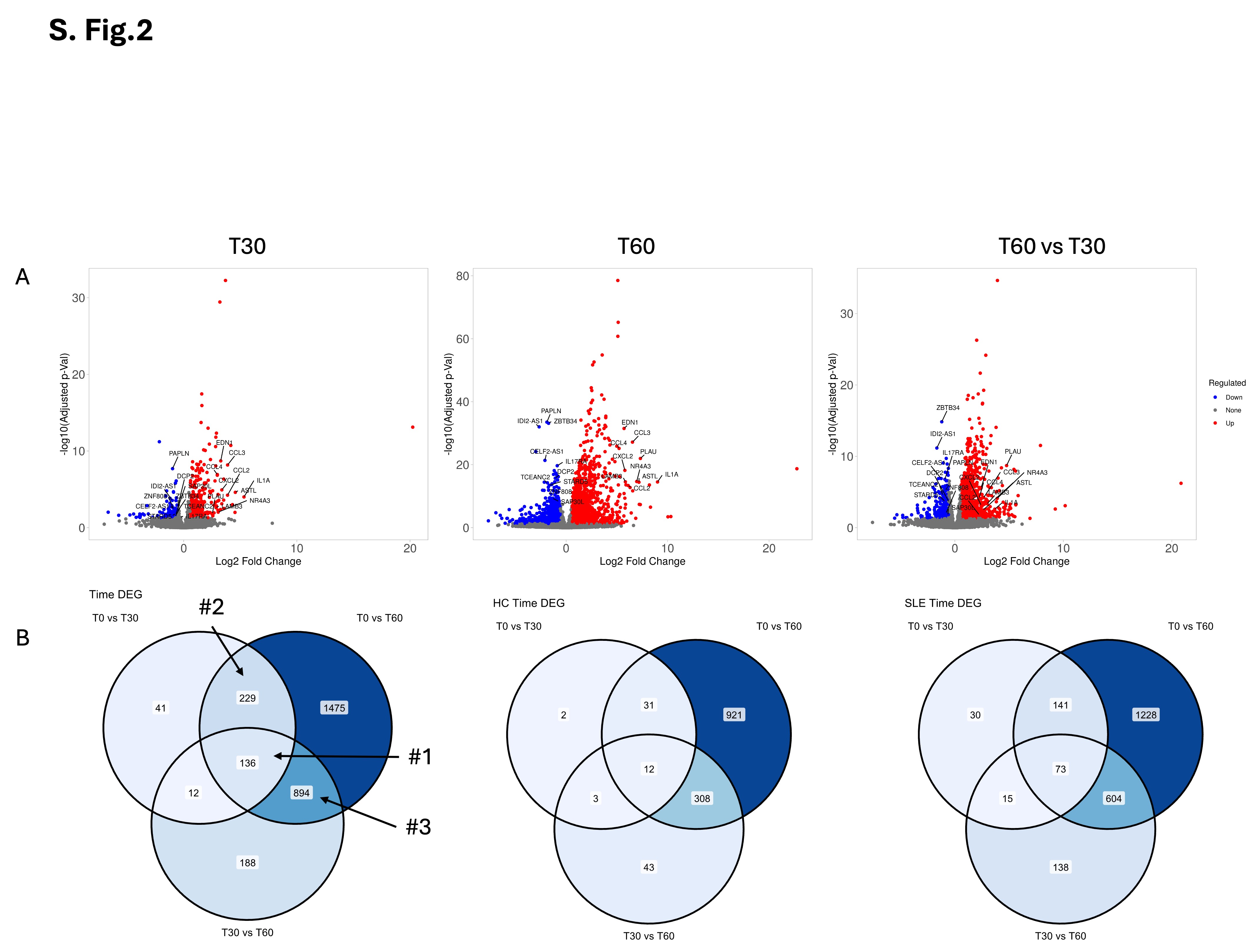

### supplementary Fig. 3

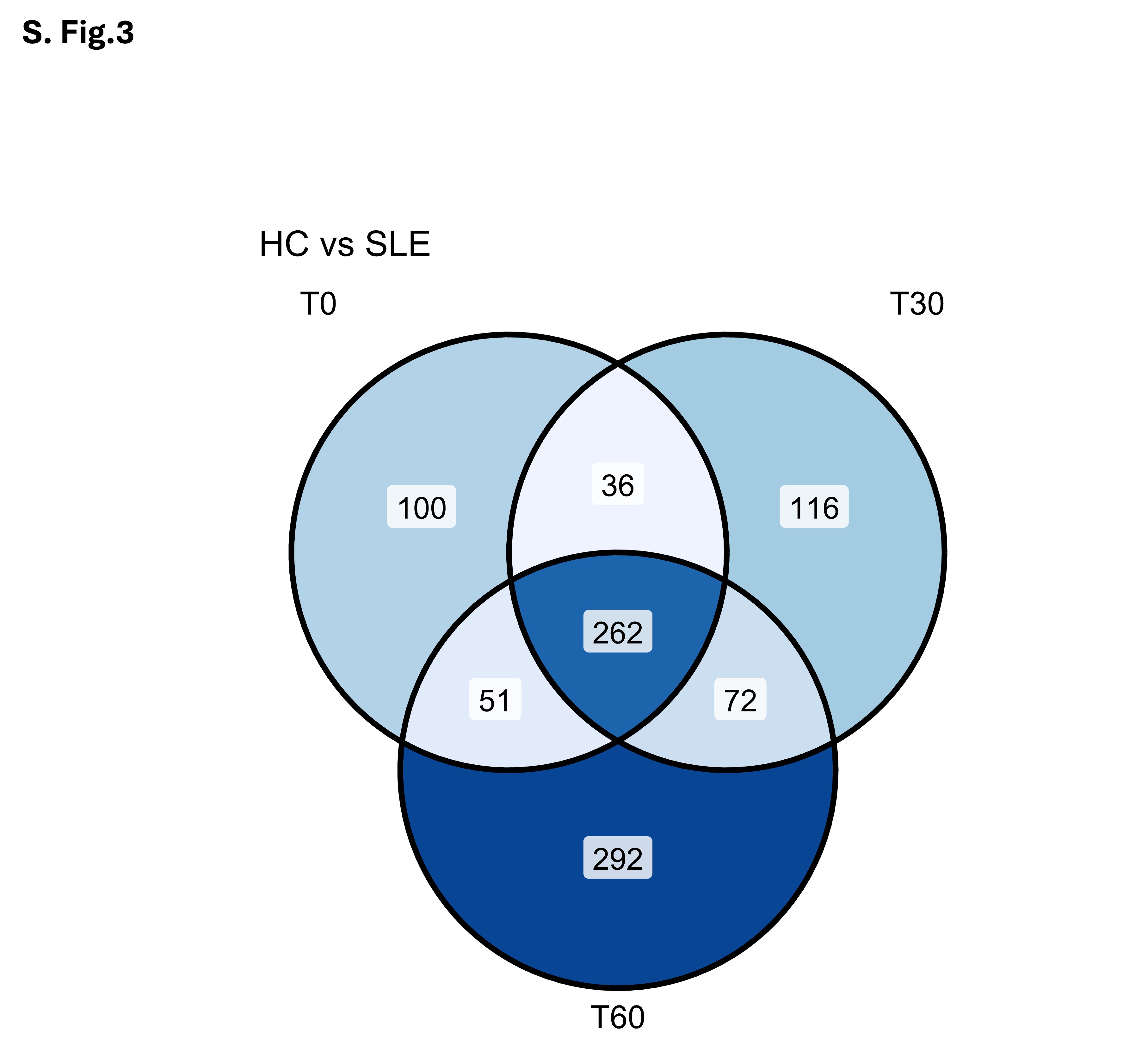

### supplementary Fig. 4

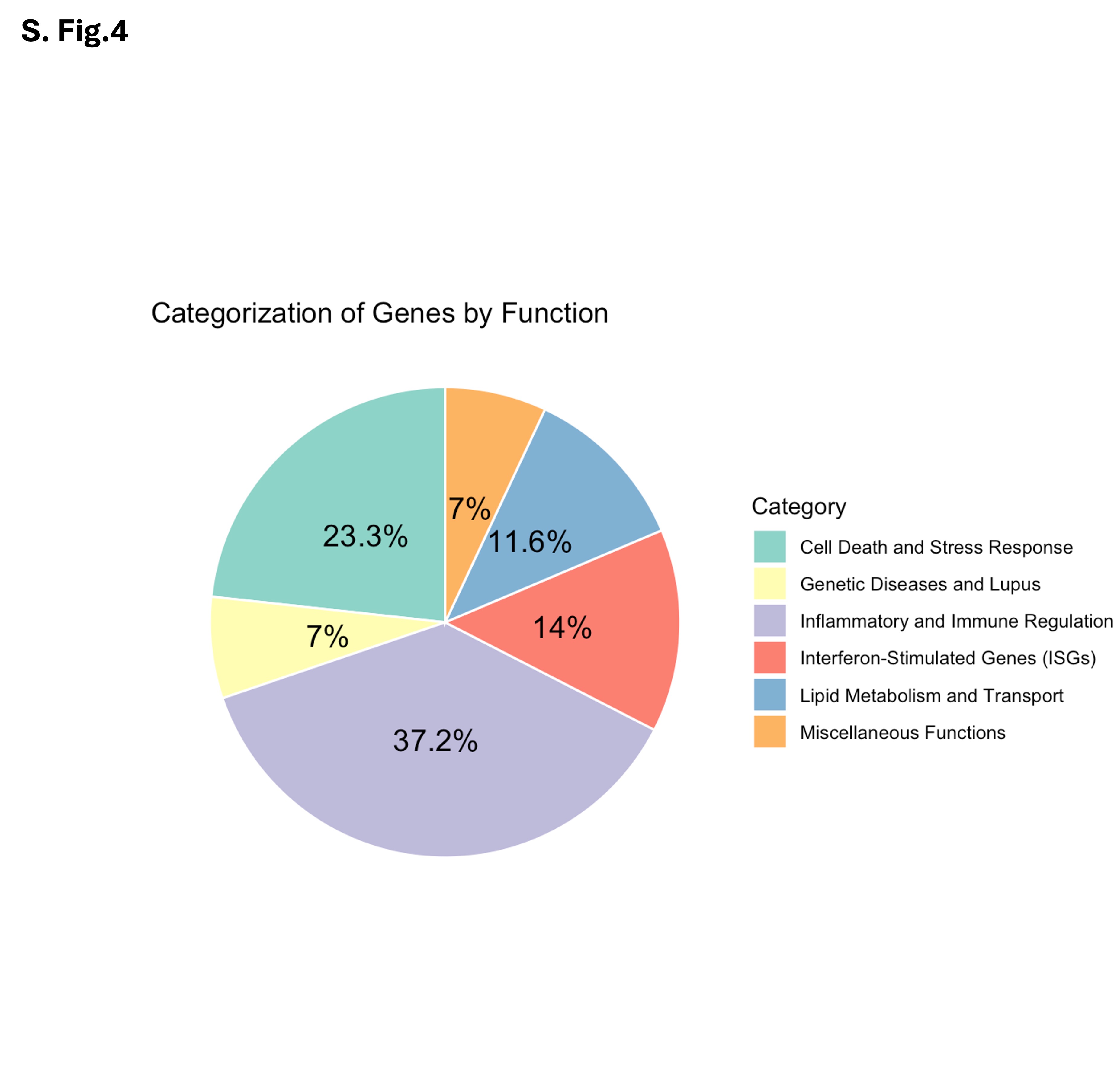

### supplementary Fig. 5

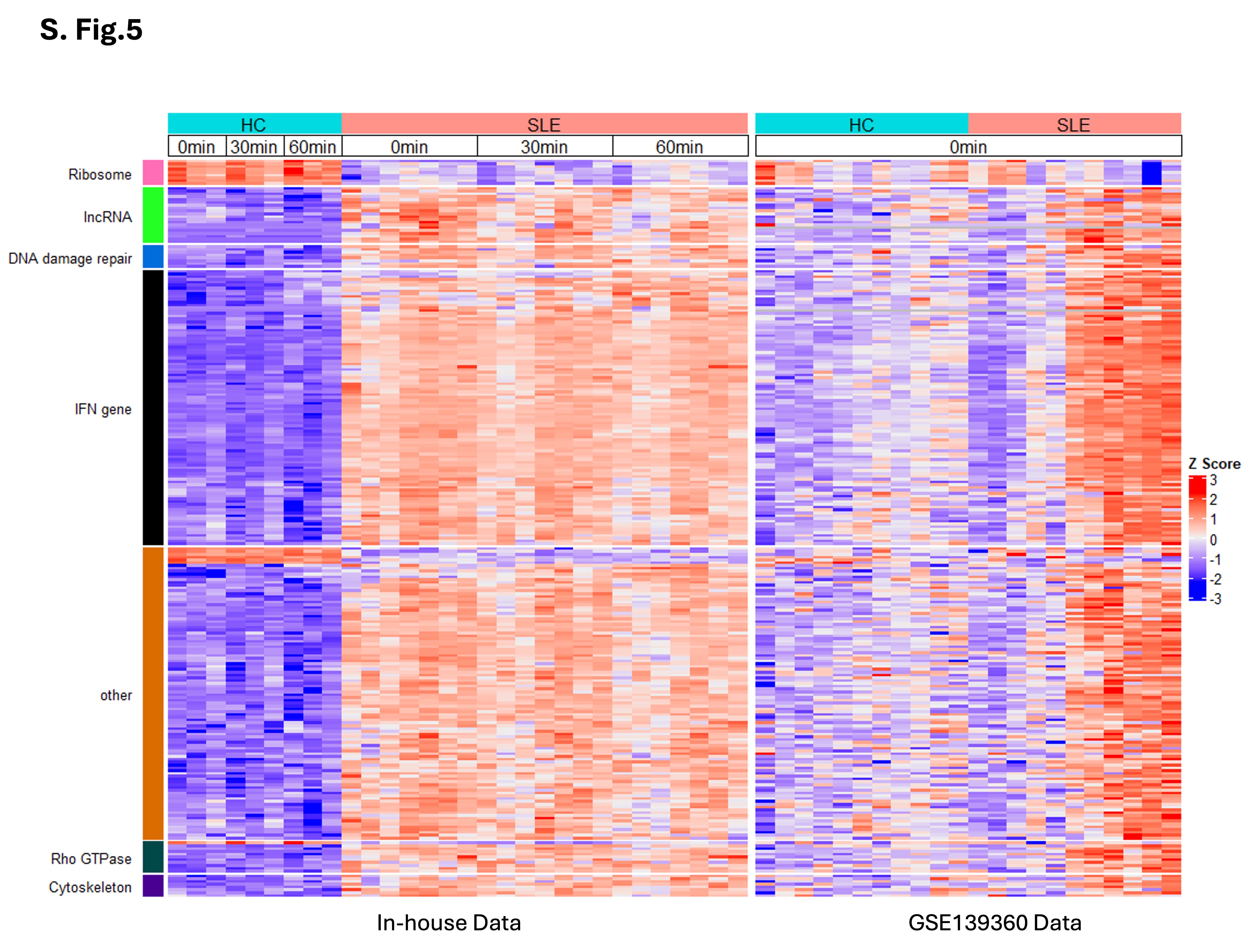

### supplementary Fig. 6

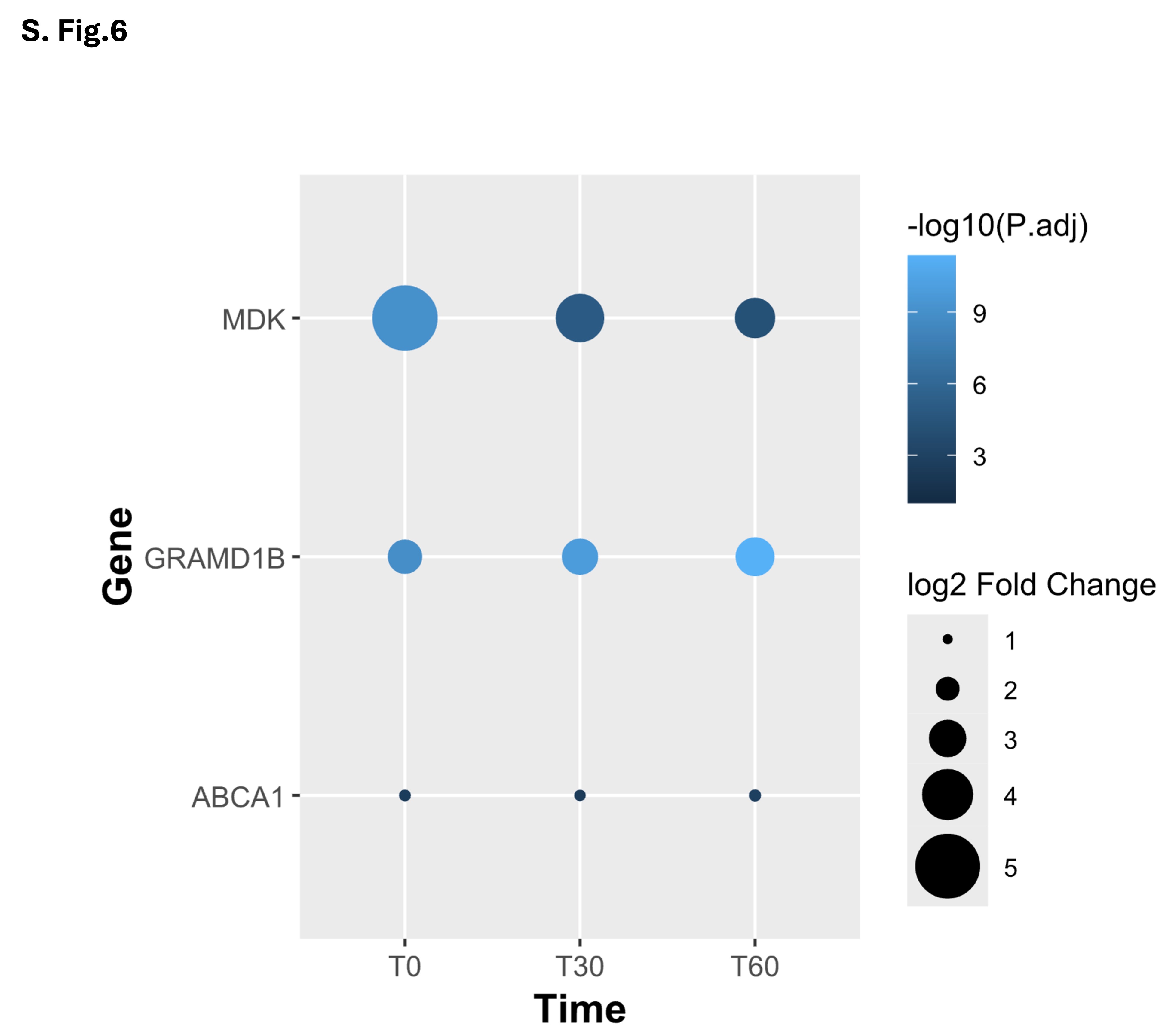

### supplementary Fig. 7

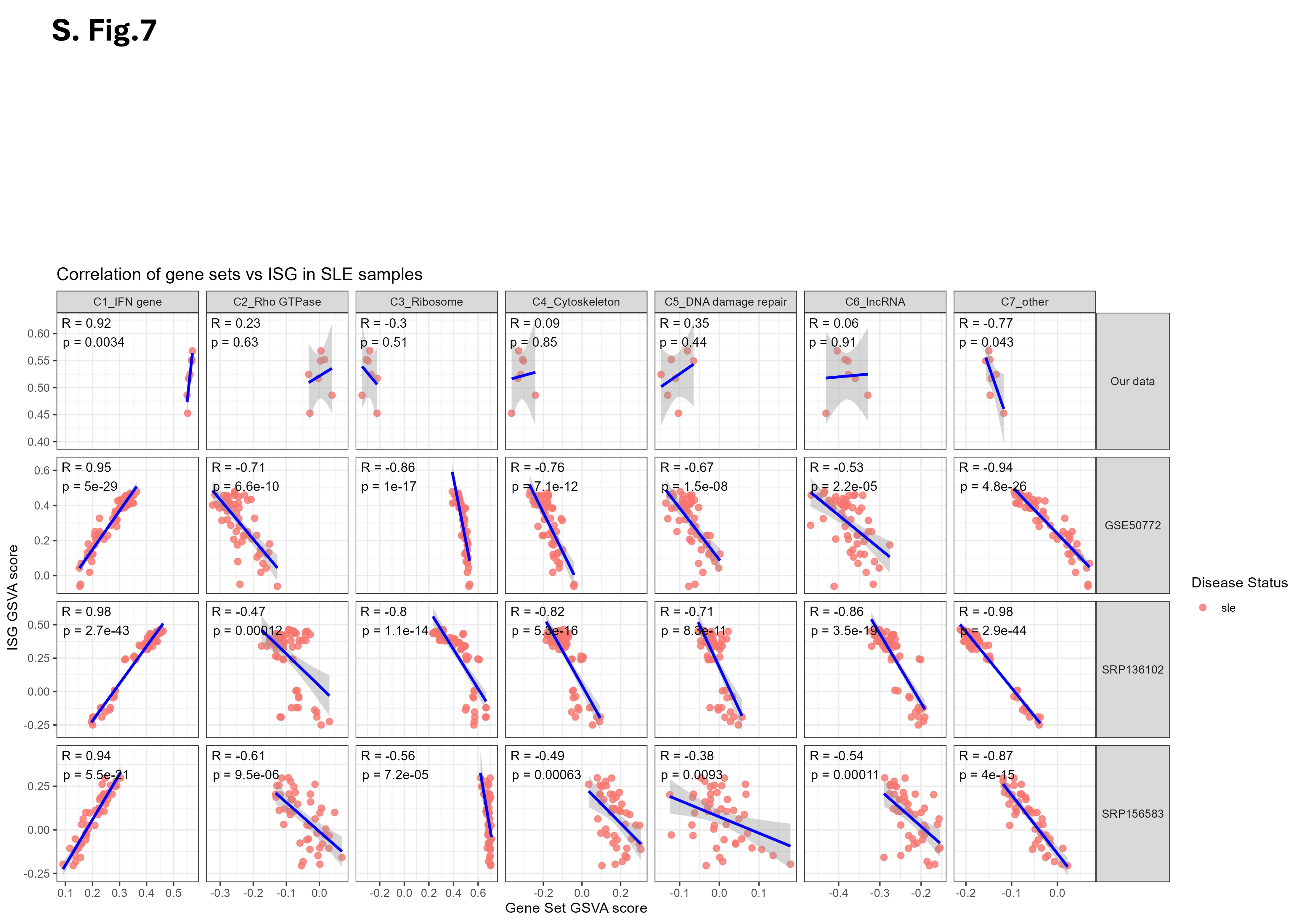
