## supplementary for "Neutrophil Transcriptomics in SLE: Exploring Intrinsic, Ex Vivo Adaptation, and CAR T-Cell Therapy-Induced Changes"

**Supplementary Figure Legends**

**Supplementary Fig. 1**

**Flow cytometric analysis of neutrophil purity following isolation.** Flow cytometry analysis of isolated neutrophils using CD45, CD16, and CD66b markers. **(A)** Representative plot showing neutrophil purity (91.7%) following isolation from buffy coat using the EasySep™ Direct Human Neutrophil Isolation Kit. **(B)** Representative plot demonstrating neutrophil purity (89.4%) after direct isolation from whole blood using the same kit. Both isolation methods yielded highly pure neutrophil populations suitable for downstream analyses.

**Supplementary Fig. 2**

**Longitudinal gene expression changes in neutrophils. (A)** Volcano plots display DEGs between time points 0 vs. 30 minutes, 0 vs. 60 minutes, and 30 vs. 60 minutes for both healthy and SLE neutrophils. **(B)** Venn diagrams showing longitudinal DEG patterns. The left diagram shows the overlap between DEGs from the three time-point comparisons (T0 vs T30, T0 vs T60, and T30 vs T60) combining both healthy and SLE samples. The middle and right diagrams display the same longitudinal comparisons separately for healthy controls and SLE patients, respectively. Despite the different numbers of DEGs, DESeq2 interaction analysis revealed no significant differences in longitudinal responses between healthy and SLE neutrophils.

**Supplementary Fig. 3**

**Venn diagram of intrinsic differentially expressed genes (DEGs).** A Venn diagram illustrates the overlap and unique gene sets between neutrophils from healthy and SLE individuals at three time points.

**Supplementary Fig. 4**

Functional categorization of "other" cluster genes. Pie chart illustrating the distribution of genes from the "other" cluster categorized by their biological functions. The largest category (37.2%) comprises genes involved in inflammatory and immune regulation, followed by cell death and stress response (23.3%), interferon-stimulated genes (14%), lipid metabolism and transport (11.6%), genetic diseases and lupus (7%), and miscellaneous functions (7%).

**Supplementary Fig. 5**

**Comparison of intrinsic DEGs across datasets.** A heatmap combining all intrinsic DEG clusters from this study, showing their expression patterns in neutrophils from healthy individuals and SLE patients. On the right, a corresponding heatmap from the GSE139360 dataset by Mistry et al. illustrates intrinsic DEG expression in normal density neutrophils from healthy and SLE individuals. The consistency between datasets highlights stable intrinsic transcriptomic differences in neutrophils from SLE patients, further validating our findings.

**Supplementary Fig. 6**

**DEGs involved in cholesterol metabolism in SLE neutrophils.** Dot plot highlighting expression of cholesterol metabolism-related genes (ABCA1, MDK, and GRAMD1B) in SLE neutrophils. Upregulation of these genes may suggest abnormalities in cholesterol handling within SLE neutrophils.

**Supplementary Fig. 7**

Correlation analysis between intrinsic DEG clusters and interferon signature genes (ISG) across multiple datasets. Scatter plots showing the correlation between GSVA scores of seven intrinsic DEG clusters (C1: IFN genes, C2: Rho GTPase, C3: Ribosome, C4: Cytoskeleton, C5: DNA damage repair, C6: lncRNA, and C7: other) and ISG scores across four datasets: neutrophils from this study (top row), PBMCs from GSE50772 (second row), whole blood from SRP136102 (third row), and B cells from SRP156583 (bottom row). Each panel displays the Pearson correlation coefficient (R) and p-value. In neutrophils from SLE patients, only the IFN-related cluster (C1) shows significant correlation with ISG (R = 0.92, p = 0.0034), while clusters C2-C6 show no significant ISG correlation. In contrast, all intrinsic DEG clusters exhibit highly significant correlations with ISG scores in PBMCs, whole blood, and B cells.

**Supplementary Table 1:**

Functional categorization and literature-supported roles of genes from the 'other' cluster. Detailed classification of genes from the "other" cluster based on their functional roles in cellular processes. Genes are grouped into six categories: Cell Death and Stress Response (including SHISA5, MICB, RNF144A, etc.), Genetic Diseases and Lupus (UNC93B1, PSTPIP2, SAT1), Inflammatory and Immune Regulation (SP140, CYSLTR1, DAPP1, etc.), Interferon-Stimulated Genes (DDX60L, SHFL, SIGLEC1, etc.), Lipid Metabolism and Transport (GRAMD1B, MDK, SLC27A3, etc.), and Miscellaneous Functions (SPATS2L, PRAL, GALM). For each gene, the table provides reference to literature supporting its functional categorization.
