## Supplementary Table 1 for "Neutrophil Transcriptomics in SLE: Exploring Intrinsic, Ex Vivo Adaptation, and CAR T-Cell Therapy-Induced Changes"

**Supplementary Table 1: Functional categorization and literature-supported roles of genes from the 'other' cluster**

| Category | Gene | Function |
| --- | --- | --- |
| **Cell Death and Stress Response** | SHISA5 | DNA damage-induced apoptosis (1) |
|  | MICB | Expression induced by DNA damage (2), tags cells for NK elimination (3) |
|  | RNF144A | Degrades PD-L1 (4), induced by DNA damage (5), promotes apoptosis (6) |
|  | TREX1 | Involved in DNA damage repair (7) |
|  | GPD2 | Inhibits apoptosis by stabilizing HIF-1α and increasing mitochondrial ROS (8) |
|  | GLRX | Anti-oxidant (9) and anti-apoptotic (10) function |
|  | EXOSC9 | Involved in cellular stress resistance (11), degradation of various RNA types (12) |
|  | CDK17 | Inhibits autophagy (13) |
|  | RUBCN | Negative regulator of canonical autophagy, positive regulator of non-canonical autophagy (14) |
|  | LGALS9 | Leads to autophagic elimination of NLRP3 in peritoneal macrophages (15) |
| **Genetic Diseases and Lupus** | UNC93B1 | Causes monogenic SLE or chilblain lupus (16, 17) (Involved in TLR trafficking) |
|  | PSTPIP2 | Loss leads to chronic multifocal osteomyelitis, increased IL-1β and ROS in neutrophils (18) |
|  | SAT1 | Loss leads to childhood-onset SLE and increased NETosis (19, 20) |
| **Inflammatory and Immune Regulation** | SP140 | Induced by IFN-γ (21), reduces LPS-induced cytokines in dendritic cells (22) |
|  | CYSLTR1 | Inhibition leads to psoriasis alleviation (23) |
|  | DAPP1 | Required for ROS production in neutrophils (24) |
|  | SCO2 | Involved in cytochrome c oxidase subunit II biogenesis (25), increases ROS production (26) |
|  | USP25 | Negative regulator of IL-17 signaling (27), decreases cholesterol biosynthesis (28) |
|  | CEACAM1 | Utilized receptor by Gram-negative bacteria that inhibits IL-1β production (29) |
|  | HSH2D | Lipoxin A4 receptor (30), role in inflammation resolution (31) |
|  | TASL | Important for IRF5 recruitment and activation in TLR 7/8/9 signaling (32) |
|  | BTN2A2 | Immunomodulatory role (33, 34) |
|  | BTN3A2 | Immunomodulatory role (35) |
|  | CCR1 | CC Chemokine receptor, involved in neutrophil chemotaxis (36) |
|  | FCMR  (downregulated in SLE) | FC mu receptor (binds specifically to IgM FC portion) (37) |
|  | PHF11 | Leads to increased IFNG transcription (38) |
|  | TFEC | Promotes M2 programming in macrophages (39) |
|  | TNFSF13B | Encodes B-cell activating factor (BAFF), activates and promotes B cell proliferation (40) |
|  | WDFY1 | Positively regulates TLR3/4-mediated signaling pathways (41) |
| **Interferon-Stimulated Genes (ISGs)** | DDX60L (42, 43), SHFL (44), SIGLEC1 (45), TMEM123 (46) | General antiviral and immune functions |
|  | ZC3HAV1 | RNA degradation during viral infection (47, 48) |
|  | NT5C3A | Induced by IFN-I (49), NF-κB pathway inhibition (50) |
|  | SLFN5 | Negative regulation of ISG expression in glioblastoma (51) |
|  | IFI16 | Induced by IFN-γ (52), intracellular DNA sensor (53) |
|  | PARP11 | Targets IFNAR1 for degradation (54) |
| **Lipid Metabolism and Transport** | GRAMD1B | Transport of cholesterol from plasma membrane to ER (55) |
|  | MDK | Inhibits cholesterol efflux in macrophages by reducing ABCA1 expression (56), elevated in SLE patients’ plasma (57) |
|  | SLC27A3 | Involved in metabolism of long-chain and very long-chain fatty acids (58) |
|  | CPT1B | Transfers long-chain fatty acids into mitochondria for β-oxidation (59, 60) (pro-inflammatory) |
|  | C7orf50 | Encodes Cholesin hormone, involved in cholesterol metabolism (61) |
| **Miscellaneous Functions** | SPATS2L | Knockdown leads to increased mRNA translation (62) |
|  | PRAL | LncRNA, increases p53 stability (63) |
|  | GALM | Involved in hexose sugars metabolism (64) |

Detailed classification of genes from the "other" cluster based on their functional roles in cellular processes. Genes are grouped into six categories: Cell Death and Stress Response (including SHISA5, MICB, RNF144A, etc.), Genetic Diseases and Lupus (UNC93B1, PSTPIP2, SAT1), Inflammatory and Immune Regulation (SP140, CYSLTR1, DAPP1, etc.), Interferon-Stimulated Genes (DDX60L, SHFL, SIGLEC1, etc.), Lipid Metabolism and Transport (GRAMD1B, MDK, SLC27A3, etc.), and Miscellaneous Functions (SPATS2L, PRAL, GALM). For each gene, the table provides reference to literature supporting its functional categorization.
